# CROP-Seq: a single-cell CRISPRi platform for characterizing candidate genes relevant to metabolic disorders in human adipocytes

**DOI:** 10.1101/2022.06.27.497796

**Authors:** Ewa Bielczyk-Maczynska, Disha Sharma, Montgomery Blencowe, Peter Saliba Gustafsson, Michael J. Gloudemans, Xia Yang, Ivan Carcamo-Orive, Martin Wabitsch, Katrin J. Svensson, Chong Y. Park, Thomas Quertermous, Joshua W. Knowles, Jiehan Li

## Abstract

**Objective:** CROP-Seq combines gene silencing using CRISPR interference (CRISPRi) with single-cell RNA sequencing (scRNA-Seq) to conduct a functional reverse genetic screen of novel gene targets associated with adipocyte differentiation or function, with single-cell transcriptomes as the readout.

**Methods:** We created a human preadipocyte SGBS cell line with stable expression of KRAB-dCas9 for CRISPRi-mediated gene knock-down. This line was transduced with a lentiviral library of sgRNAs targeting 6 genes of interest (3 sgRNAs / gene, 18 sgRNAs), 6 positive control genes (3 sgRNAs / gene, 18 sgRNAs), and non-targeting control sgRNAs (4 sgRNAs). Transduced cells were selected and differentiated, and individual cells were captured using microfluidics at day 0, 4 and 8 of adipogenic differentiation. Next, expression and sgRNA libraries were created and sequenced. Bioinformatic analysis of resulting scRNA-Seq expression data was used to determine the effects of gene knock-down and the dysregulated pathways, and to predict cellular phenotypes.

**Results:** Single-cell transcriptomes obtained from SGBS cells following CRISPRi recapitulate different states of differentiation from preadipocytes to adipocytes. We confirmed successful knock-down of targeted genes. Transcriptome-wide changes were observed for all targeted genes, with over 400 differentially expressed genes identified per gene at least at one timepoint. Knock-down of known adipogenesis regulators *PPARG* and *CEBPB* inhibited adipogenesis. Gene set enrichment analyses revealed molecular processes for adipose tissue differentiation and function for novel genes. *MAFF* knock-down led to a downregulation of transcriptional response to proinflammatory cytokine TNF-α in preadipocytes. *TIPARP* knock-down resulted in an increase in the expression of a beiging marker *UCP1* at D8 of adipogenesis.

**Conclusions:** The CROP-Seq system in SGBS cells can determine the consequences of target gene knock-down at the transcriptome level. This powerful, hypothesis-free tool can identify novel regulators of adipogenesis, preadipocyte and adipocyte function associated with metabolic disease.

**Highlights:**

- CRISPR interference screen coupled with single-cell RNA sequencing (CROP-Seq)
- Parallel screening of 12 genes in human SGBS adipocytes and preadipocytes
- Uncovered novel regulators of adipogenesis and adipocyte function

**Graphical abstract:** 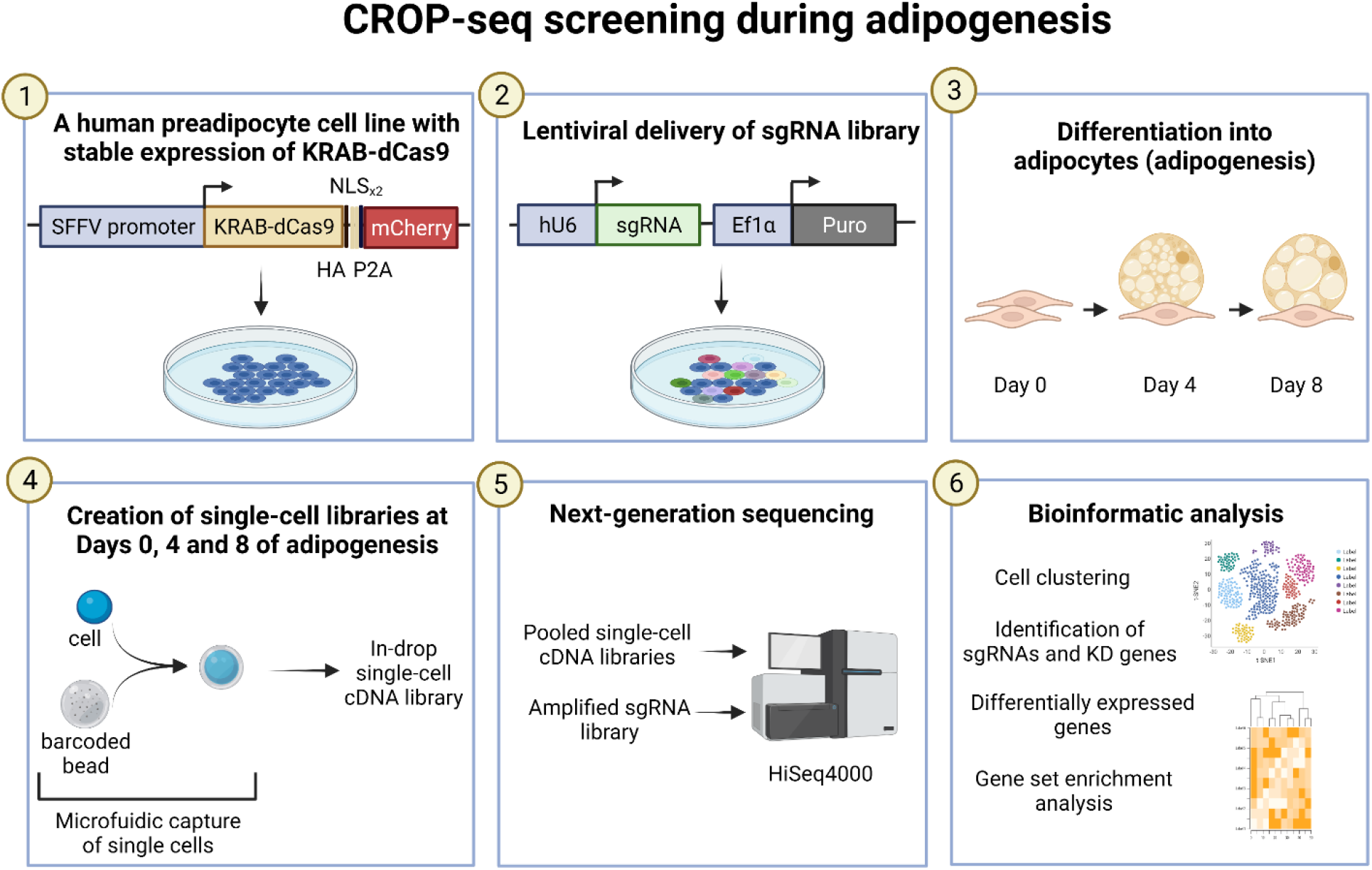

## 1. Introduction

Over the past decade, numerous genomics efforts uncovered a wealth of associations between human genetic loci and metabolic traits, creating a rich resource that can be mined to identify novel factors contributing to disease pathophysiology. However, within many of these genetic loci, the causal gene remains unclear hampering downstream investigations^1^. Many of these trait loci are tied to adipose tissue which is not surprising given the key role of adipose tissue in metabolic diseases. Differences in both adipocyte differentiation and adipocyte function contribute to metabolic disorders. For example, in lipodystrophies insufficient adipocyte differentiation from preadipocyte progenitors leads to insulin resistance and type 2 diabetes (T2D)^2^. Adipocyte hypertrophy contributes to metabolic abnormalities associated with obesity^3^.

Additionally, adipocyte browning/beiging improves whole-body metabolism in rodent models^4^ and is associated with leanness in humans^5,6^. As diverse adipocyte-related phenotypes can contribute to whole-body metabolic health, there is an advantage to using hypothesis-free approaches to investigate the molecular role of candidate genes within disease-associated loci. Once a potential molecular mechanism is identified, gene function can be validated using a relevant assay.

Functional follow-up of novel genes predicted to affect metabolism through their adipocyte-related function is a time-consuming process. Creation of knock-out mouse lines is costly and low-throughput. In addition, there are key differences between some metabolically-relevant processes in humans and mice, for example in cholesterol metabolism^7^. In cell culture models, prolonged culture of individual genetically manipulated lines may lead to artifacts resulting from clonal effects. Therefore, pooled screens that can be accomplished within a short time frame, in which different experimental groups and controls are subjected to the same handling and environmental stimuli, are advantageous.

Here, we establish a platform for high-throughput studies of genes involved in adipogenesis and adipocyte function in a human non-immortalized Simpson-Golabi-Behmel syndrome (SGBS) model^8^. The CROP-Seq platform combines CRISPRi-mediated pooled gene knock-down with single-cell RNA sequencing (scRNA-Seq) using DROP-seq^9^ to determine transcriptomic states resulting from target gene knock-down. As a proof-of-concept, we show that this approach captures the phenotypes resulting from knock-down of known regulators of adipogenesis. Further, we show that it can be utilized to generate hypotheses regarding the role of novel genes in distinct mechanisms relevant for adipose tissue function.

## 2. Materials and methods

### 2.1 Gene selection

Candidate genes were selected in 2019 based on colocalization of GWAS hits for metabolic traits: waist-hip ratio adjusted for BMI (WHRadjBMI)^10^, fasting insulin (FI)^11^, fasting glucose (FG)^11^, triglycerides (TG)^12^, HDL-cholesterol^12^ with eQTL in adipose tissue (subcutaneous or visceral) as described^13^. The gene list was further limited to genes with Unique Molecular Identifier (UMI) expression of at least 2 in at least 9% of individual cells at day 7 (D7) of adipogenesis, based on a scRNA-Seq of SGBS cells (data not shown).

### 2.2 Cell culture and differentiation

The SGBS preadipocyte cell line was cultured and differentiated as described before^8^. Cells were maintained in a humidified chamber at 37°C with 5% CO_2_, and their media was replaced every 2-3 days. The standard culture media used was composed of DMEM/Nutrient Mix F-12 (Invitrogen), supplemented with 33 uM biotin, 17 uM pantothenic acid, 10 % FBS and antibiotics (100 IU/ml penicillin and 100 ug/ml streptomycin). Differentiation was induced three days post-confluence by the change of culture media to DMEM/F-12, 33 Um biotin, 17 uM pantothenic acid, 0.01 mg/ml human transferrin, 100 nM cortisol, 200 pM triiodothyronine, 20 nM human insulin (Sigma-Aldrich), 25 nM dexamethasone, 250 uM IBMX, 2 uM rosiglitazone, and antibiotics. After four days of differentiation, the medium was replaced with DMEM/F-12, 33 uM biotin, 17 uM pantothenic acid, 0.01 mg/ml human transferrin, 100 nM cortisol, 200 pM triiodothyronine, 20 nM human insulin and antibiotics. Cells were cultured for up to eight days after the induction of differentiation. As differentiation ability of SGBS cells is affected by local confluence, and lentiviral transduction and subsequent puromycin selection may affect their proliferation ability, cells were plated evenly at high density.

To validate the transcriptional effects of TIPARP knock-down, the TIPARP inhibitor RBN-2397 (Fisher Scientific) was used at a concentration of 1 uM. Recombinant human TNF-alpha (PeproTech/VWR) was used at a concentration of 10 ng/ml.

### 2.3 Quantitative PCR

Total RNA was isolated from cells using the RNeasy Mini kit (QIAGEN) according to the manufacturer’s instructions. Reverse transcription was conducted using High Capacity cDNA Reverse Transcription Kit (Thermo Fisher Scientific). Gene expression was quantified on the ViiA7 machine (Applied Biosystems), using Taqman Gene Expression Master Mix (Applied Biosystems) and standard cycling settings. The following gene expression assays were obtained from Thermo Fisher Scientific: *CXCL6* (Hs00605742_g1), *IL6* (Hs.PT.58.39866843.g), *PPARG* (Hs01115513_m1), *PPIA* (Hs04194521_s1), *UCP1* (Hs01084772_m1). The following PrimeTime qPCR Probe Arrays were obtained from Integrated DNA Technologies: *CYP1B1* (Hs.PT.58.25328727.g), *STC2* (Hs.PT.58.38362090). All gene expression results were normalized to the expression of the housekeeping gene *PPIA* using the ddCt method.

### 2.4 siRNA-mediated gene knock-down

siRNA duplexes targeting *TIPARP* and Universal scrambled negative control were purchased from Origene and transfected into SGBS cells using Lipofectamine RNAiMAX reagent (Invitrogen) according to the manufacturer’s protocol.

### 2.5 Generation and validation of the stable SGBS CRISPRi cell line

pHR-SFFV-KRAB-dCas9-P2A-mCherry plasmid (Addgene, #60954) was a gift from Jonathan Weissman^14^. Lentivirus was produced as previously described^15^. Transduced cells were purified based on mCherry expression using FACS. The resulting cell line was validated by transduction with sgRNAs targeting individual genes, cloned into the pCRISPRia-v2 plasmid digested with BstXI and BlpI (New England Biolabs). Following the sgRNA transduction and puromycin selection, gene expression prior to differentiation (D0) and adipogenic marker gene expression at D8 of differentiation was quantified using RT-qPCR.

### 2.6 sgRNA cloning and CROP-Seq plasmid library preparation

Three sgRNAs targeting each gene of interest, and four control non-targeting sgRNAs, were chosen from the human genome-wide CRISPRi-v2 library^16^ **(Supplementary Table 1)**. The inserts were ordered as ssDNA from Integrated DNA Technologies. They consisted of the sgRNA sequence with the following overhangs: TGGAAAGGACGAAACACCG (5’) and GTTTTAGAGCTAGAAATAGCAAGTTAAAATAAGGC (3’). The CROP-seq-opti plasmid (Addgene, #106280) was digested with BsmBI and purified from agarose gel using QIAquick Gel Extraction Kit (QIAGEN). Ligation was conducted using NEBuilder HiFi DNA Assembly Master Mix (New England Biolabs) and transfected into NEB Stable *E*.*coli*. Single colonies were picked and the correct sequence of the inserts was validated using Sanger sequencing. The plasmids were combined at equal ratios into a lentiviral plasmid library for the CROP-Seq screening. For validation experiments some of the plasmids were also used individually.

### 2.8 Cell transduction, selection and differentiation

Lentiviral stock was produced as described previously^15^. To establish the dilution of the lentiviral stock that results in multiplicity of infection (MOI) of 0.3-0.4, SGBS cells were transduced with various lentivirus titers, cultured for two days, selected with puromycin (1 ug/ml) for two days, and allowed to recover in puromycin-free media for one day. Next, cell number was approximated using alamarBlue Cell Viability Reagent (Thermo Fisher Scientific) according to the manufacturer’s protocol.

In the CROP-Seq experiment, SGBS cells were transduced with the selected lentiviral dilution associated with MOI 0.3-0.4, cultured for two days, and selected with puromycin for two days, followed by one-day recovery in puromycin-free media. Next, cells were replated at 5,000 cells / cm^2^ in 12-well plates and cultured for further 3-5 days until confluence. Once cells were confluent, standard differentiation protocol was followed. At D0, D4 and D8 of differentiation, cells were trypsinized and subjected to cell sorting.

### 2.9 Cell sorting and CROP-Seq library construction and sequencing

On the day of the single-cell capture, cells were trypsinized, filtered and stained with the live/dead stain SYTOX Green Ready Flow Reagent (Thermo Fisher) according to the manufacturer’s protocol. Cells were stored on ice throughout the procedure. Live (SYTOX-negative) mCherry-positive single cells were isolated using fluorescence-activated cell sorting (FACS) at the Stanford Shared FACS Facility. Next, live cells underwent microfluidic single-cell capture on the 10X Chromium Controller device at Stanford Genomics Service Center during which single cells were encapsulated with individual Gel Beads-in-emulsion (GEMs) using the Chromium Next GEM Single Cell 3’ GEM, Library & Gel Bead Kit (10X Genomics). In-drop reverse transcription and cDNA amplification was conducted according to the manufacturer’s protocol to construct expression libraries. Library size was checked using Agilent Bioanalyzer 2100 at the Stanford Genomics facility. Expression libraries were sequenced by Novogene on the HiSeq4000 sequencer.

Additional sequencing was performed to determine sgRNA representation and to assign sgRNA sequences to individual cells. To determine sgRNA representation, sgRNA sequences were amplified from bulk genomic DNA after transduction, at D0 and D8 of differentiation. To assign sgRNAs to individual cell barcodes, sgRNA sequences were amplified from the pooled cDNA. PCR products were purified using AMPure XP (Beckman Coulter) and the SPRIselect reagent (Beckman Coulter) according to the manufacturer’s protocol. All sgRNA libraries were combined and sequenced on the HiSeq4000 instrument by Novogene.

### 2.10 Sequencing data analysis

The sequencing data from SGBS cells in the format of FASTQ files were aligned to the human (hg38) reference genome and quantified using “cellranger count” (10X Genomics). The aligned files were then analyzed using R package Seurat. The dataset was trimmed of cells expressing fewer than 200 genes. Cells with mitochondria content higher than 5% were considered poor quality cells and were discarded from the dataset. The gene expression matrix was then used to perform normalization and scaling using Seurat’s built-in functions. Principal component analysis for dimensionality reduction followed by clustering using graph-based clustering approach was performed. The UMAP was then used to visualize the resulting clusters in the two-dimensional space. “FeaturePlot” and “DotPlot” functions from Seurat’s pipeline were used to visualize the expression of the gene of interest. We used MAST to identify differentially expressed (DE) genes with p-value < 0.05 and log2FC cut-off 0.5.

Lineage inference (pseudotime analysis) was performed using the Dyno package which includes a collection of pseudotime and trajectory inference tools. We performed trajectory inference with expected topology as “bifurcating” so the best model was built using “slingshot”, “page_tree”, “paga” and “mst”.

Due to high percentage of multiple gRNA cells, we used scMAGeCK pipeline^17^ for single-cell CRISPR screening data. After filtering against the significance as well as degree of perturbation impact, we ranked all the target genes by the number of differentially expressed genes.

### 2.11 Gene set enrichment analysis

Gene set enrichment analysis was conducted using the Hallmark gene sets^18^ and Gene Ontology^19,20^ gene sets within the MsigDB collections^21^. Genes identified by scMAGeCK with adjusted p-value < 0.05 were used and scMAGeCK score cut-off at <(−0.1) or >0.1. Up to the top 500 DE genes from scMAGeCK, ranked by the p-value, were used for the analysis. All the reported pathways were shown with the corresponding q-values.

### 2.12 Co-expression module build and gene query

Human co-expression networks were constructed from transcriptome data from adipose tissue samples, including visceral omentum and subcutaneous adipose from the GTEx study^22^. Mouse co-expression networks were constructed from transcriptome data from adipose tissue from 101 Hybrid Mouse Diversity Panel (HMDP) strains^23^, which included healthy mice, mice with liver steatosis and mice with nonalcoholic steatohepatitis (NASH). The co-expression networks were built using two network methods: Weighted Gene Co-expression Network Analysis (WGCNA)^24^ and Multiscale Embedded Gene Co-expression Network Analysis (MEGENA)^25^. WGCNA and MEGENA cluster genes into modules based on the co-regulation structure of the genes using hierarchical clustering. Modules from both methods can be reciprocally conserved, however, their compactness and sizes are different, making these two methods complementary. Once these modules were built, they were queried for candidate genes to identify co-expressed genes. For any module that contained either a candidate or control gene, we annotated it with its functions by using the previously curated biological pathways taken from MSigDB database^26^ that incorporates pathways from Biocarta, KEGG, and Reactome databases based on a hypergeometric test (one-tailed Fisher Exact test). Bonferroni correction was used to adjust the p-values of this test. Pathways with a corrected p-value<0.05 and sharing ≥ 5 genes with a given module were accepted as significant. The top three significant pathways were used to annotate each module.

### 2.13 Statistical analyses

For functional assays, normal distribution of data was tested using Kolmogorov-Smirnov test. Differences between groups were compared using one-way ANOVA, with p-value < 0.05 interpreted as significant. Tukey’s multiple comparisons test was used to correct for multiple testing. Unless indicated otherwise, average and SEM are shown. N=3 biological replicates were used unless indicated otherwise. Statistical analyses and plotting were conducted using GraphPad Prism 9.

## 3. Results

### 3.1 Creation of the CROP-Seq platform in the human preadipocyte SGBS cell line

In the CROP-Seq experiment, a human preadipocyte SGBS cell line with stable expression of KRAB-dCas9 was transduced with a pooled lentiviral library containing sgRNAs targeting the genes of interest and non-targeting control sgRNAs. The transduced cells were sorted and subjected to scRNA-Seq using DROP-Seq either before (Day 0) or during (Day 4 and Day 8) differentiation, followed by next-generation sequencing and bioinformatic analysis.

To establish the CRISPRi SGBS line, we introduced KRAB-dCas9-P2A-mCherry transgene into wild-type SGBS cells using lentivirus, followed by FACS-based selection of mCherry-expressing cells. Next, to validate that the cell line contains functional CRISPRi machinery, SGBS CRISPRi cells were transduced with lentivirus encoding sgRNAs targeting transcription factors which drive adipogenesis *(CEBPB* and *PPARG*) or non-targeting sgRNAs as a control. Robust downregulation of target gene expression at D0 and D8 of differentiation **(Supplementary Figure 1A,B)**, as well as predicted downregulation of the adipogenesis marker *FABP4* at D8 by over 60% **(Supplementary Figure 1C)**, was confirmed using RT-qPCR.

To select candidate genes for the screen, we used a colocalization approach^13^, which combines information from genome-wide analysis studies (GWAS) with expression quantitative trait loci (eQTLs). We focused on GWAS relevant for metabolic disease traits: WHRadjBMI^27^, fasting insulin^11^, fasting glucose^11^, triglycerides^12^, and HDL-cholesterol^12^. eQTLs for subcutaneous (SubQ) and visceral adipose tissue were selected. For the colocalization analysis, genes with a minimum colocalization posterior probability (CLPP) score^28^ of 30% for at least one of the traits colocalizing with adipose tissue were chosen. Candidate genes were further selected based on their expression level in single SGBS cells (unpublished data) and literature search indicating their novelty. The following candidate genes were chosen: *FAM13A, MAFF, PDGFC, TIPARP, VEGFB*, and *ZBTB7B*. In addition, positive control genes were included based on their known role in adipogenesis (*PPARG*^29^, *CEBPB*^30^), insulin signaling (*IRS1*^31^), lipid metabolism (*MLXIPL*^32^), or their role as adipokine-encoding genes (*ADIPOQ*^33^, *RBP4*^*34*^). The pooled lentiviral library was composed of three sgRNAs per gene, as well as 10% non-targeting sgRNAs which served as negative controls **(Supplementary Table 1)**. The pooled screen was conducted, and single cells were collected at D0, D4 and D8 of adipogenesis.

To confirm whether sgRNAs affect the expression of their target genes, we compared average expression and percentage of cells with detectable expression of candidate gene depending on the sgRNA that was present. We observed noticeable downregulation of the target transcripts for most sgRNAs at all three time points **(Figure 1A, Supplementary Figure 1D)**. Altogether, we established that the CROP-Seq approach captures single-cell transcriptome data, with concurrent target gene repression. To our knowledge, this is the first study which used this type of single-cell perturbation and readout to study adipogenesis using the human SGBS cell model.

**Figure 1:**
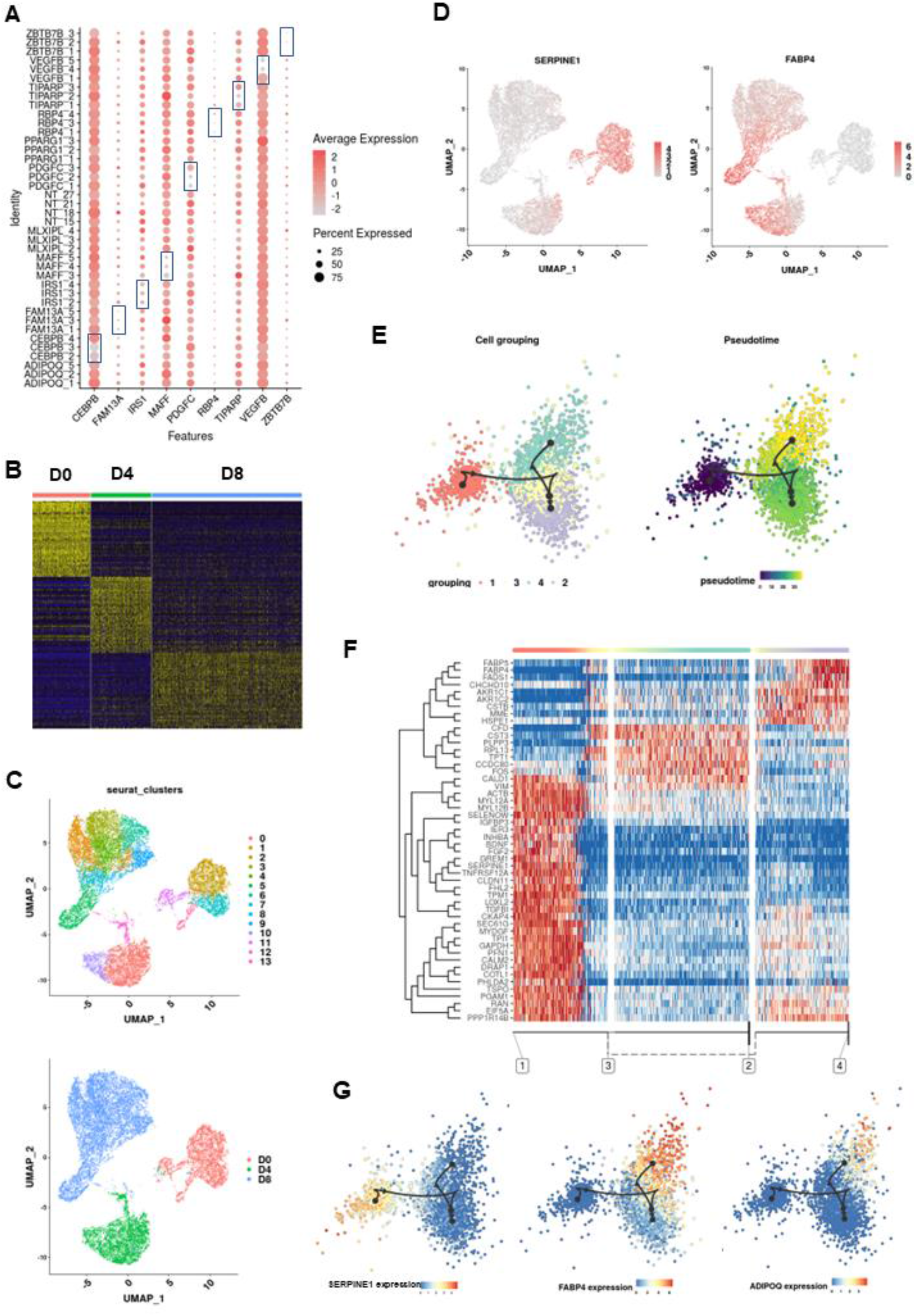
Establishment of a CRISPRi screen in a human primary cell model of adipocyte differentiation at single-cell resolution. **A)** Quantification of target gene expression at D0 by average expression and percent of cells with detectable expression depending on the sgRNA. knock-down of targeted genes is highlighted by the blue boxes. **B)** Heat map of all differentially expressed (DE) genes within cells with non-targeting sgRNAs using Seurat shows clustering by day of differentiation. **C)** UMAP plot of all cells with identified sgRNAs across differentiation timepoints. **D)** UMAP plot of all cells showing single-cell expression of marker genes for pre-adipocytes (*SERPINE1*) and adipocytes (*FABP4*). **E)** Slingshot pseudotime ordering of all non-targeting sgRNA-expressing cells reveals a continuous trajectory of cell states corresponding to adipocyte differentiation. **F)** Heatmap of genes driving Slingshot pseudotime states in cells with non-targeting sgRNAs. **G)** Expression of preadipocyte (*SERPINE1*) and adipocyte (*ADIPOQ, FABP4*) markers along the pseudotime trajectory.

To determine whether cells within the CROP-Seq experiment recapitulate transcriptional states expected for differentiating preadipocytes, we first compared transcriptional states of cells with non-targeting sgRNAs only. Clustering of differentially expressed (DE) genes revealed a clear separation of differentiation states at D0, D4 and D8 **(Figure 1B)**. Moreover, unsupervised clustering of all cells with assigned sgRNAs revealed 10 cell clusters, with three larger clusters corresponding to the day of differentiation **(Figure 1C)**. Cells with non-targeting sgRNAs were represented in all clusters **(Supplementary Figure 1E)**. The three clusters showed the predicted pattern of expression of marker genes for preadipocytes (*SERPINE1*^35^, *GREM1*^36^) and adipocytes (*ADIPOQ*^*37*^, *FABP4*^*38*^)**(Figure 1D, Supplementary Figure 1F)**.

### 3.2 CROP-Seq identifies transcriptional states along the preadipocyte to adipocyte differentiation trajectory

Pseudotime trajectory analysis of cells with non-targeting sgRNAs was used to infer cell lineages, which revealed cells grouping in four main states **(Figure 1E)**. Analysis of the heatmap containing the genes which drove the pseudotime trajectory analysis showed that cells clustered largely based on the day of differentiation **(Figure 1F)**. Notably, cells collected at D8 and D4 showed a distribution of transcriptional states, likely due to a mixture of undifferentiated, partially differentiated and differentiated cells **(Figure 1F)**. Analysis of single-cell expression of adipocyte (*ADIPOQ, FABP4*) and preadipocyte (*SERPINE1*) marker genes further supported that the pseudotime trajectory replicates the predicted states during adipogenic differentiation **(Figure 1G)**. Altogether, these complementary approaches showed that the transcriptional states captured in the CROP-Seq experiment are consistent with a mixture of preadipocytes, differentiating cells and adipocytes.

### 3.3 Knock-down of *PPARG* and *CEBPB* results in adipogenesis inhibition

Next, to establish analysis approach that would allow us to identify gene networks whose expression is altered following the knock-down of candidate genes, we focused on genes which were differentially expressed (DE) following *PPARG* knock-down. PPARγ is a key adipogenic transcription factor whose pattern of gene expression regulation in SGBS adipocytes is well-described^39-41^. PPARγ binds to the promoters of over 5,600 human genes, with approximately 2,700 genes (corresponding to 10-14% of all human genes) showing conserved binding between human and mouse adipocytes^41^. Based on these data, we used additional filtering based on p-value and score to select DE genes resulting from scMAGeCK analysis^17^ **(Supplementary Figure 2)**. The number of DE genes varied across the candidate genes targeted and differentiation timepoints, with the highest average DE gene number for *PPARG* and *CEBPB* knock-down (score <(−0.1) or >0.1 and p-value < 0.05) **(Figure 2A)**. For *PPARG* knock-down, the number of DE genes was the highest at D8. Expectedly, Gene Set Enrichment Analysis (GSEA) showed a significant enrichment of genes involved in adipogenesis within the *PPARG* DE genes at D4 and D8 **(Figure 2B)**, with many known PPARG targets amongst the downregulated DE genes **(Figure 2C)**. Altogether, these data supported differentiation inhibition in *PPARG* knock-down cells^29,42^.

**Figure 2:**
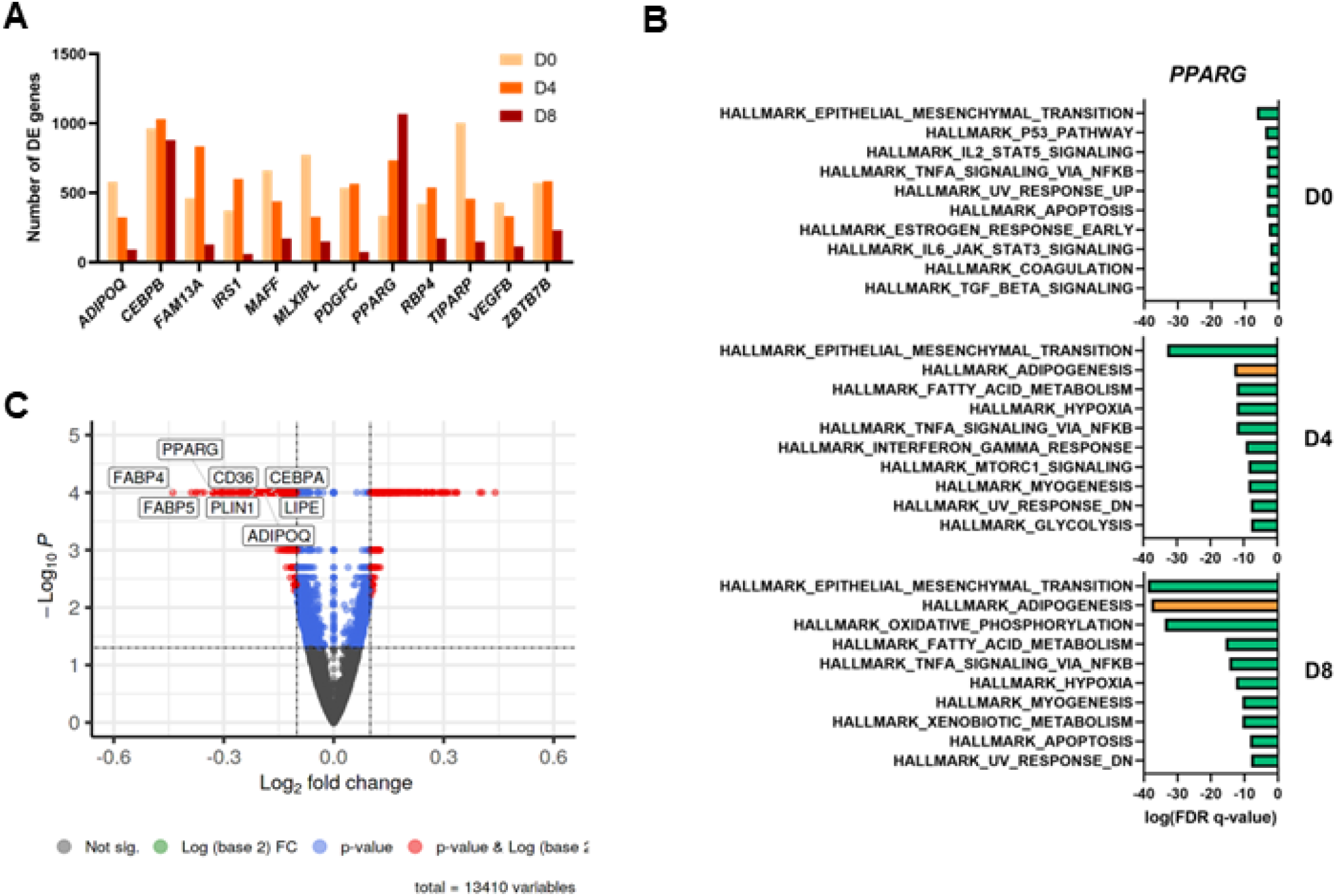
CROP-seq is a reliable approach to characterize biologically meaningful genetic perturbation effects in the human SGBS adipogenesis model. **A)** Number of differentially expressed genes identified using scMaGeCK in different gene knock-down conditions with score 0.1 cutoff and p < 0.05. **B)** GSEA analysis of *PPARG* knock-down DE genes at D0, D4, D8. Gene set of adipogenesis-related genes is highlighted in orange. **C)** Volcano plot of DE genes for *PPARG* knock-down at D8.

Similar to PPAR-γ, C/EBP-β is an adipogenic transcription factor. However, it acts at an earlier stage of adipogenesis compared to PPAR-γ^43,44^. Consistent with the known role of C/EBP-β in early adipogenesis, the highest number of DE for *CEBPB* knock-down were identified at D4 and D0 **(Figure 2A)**. GSEA analysis pointed to changes in the expression of adipogenesis-related genes at D4 and D8, but not D0 **(Supplementary Figure 3)**. In conclusion, knock-down of genes encoding known regulators of adipogenesis, *PPARG* and *CEBPB*, was used as a proof-of-concept for the use of the CROP-Seq identify gene expression patterns affected by candidate gene downregulation.

### 3.4 *MAFF* regulates cellular preadipocyte response to TNF-α

To determine whether CROP-Seq can be used as a discovery tool to identify novel gene function, we focused on one of the candidate genes, *MAFF*, which encodes the MAF Basic Leucine Zipper Transcription Factor F. *MAFF* was identified in GWAS on HDL and triglyceride levels^45^. *MAFF* variants have also been identified as associated with T2D risk^46^. *MAFF* eQTL signal colocalized in both subcutaneous and visceral adipose tissue. However, its function in adipose tissue biology has not been studied to date.

*MAFF* knock-down led to largest changes in gene expression at D0, assessed by the number of DE genes **(Figure 2A)**. Gene Set Enrichment analysis revealed that *MAFF* knock-down at D0 was associated with changes in the expression of genes related to TNF-α signaling via the NF-κB pathway **(Figure 3A)**. In support of the possible role of MAFF in mediating cellular response to TNF-α, co-expression analyses of *MAFF* in human adipose tissue consistently indicated *MAFF* involvement in cytokine signaling-related networks **(Figure 3B)**.

**Figure 3:**
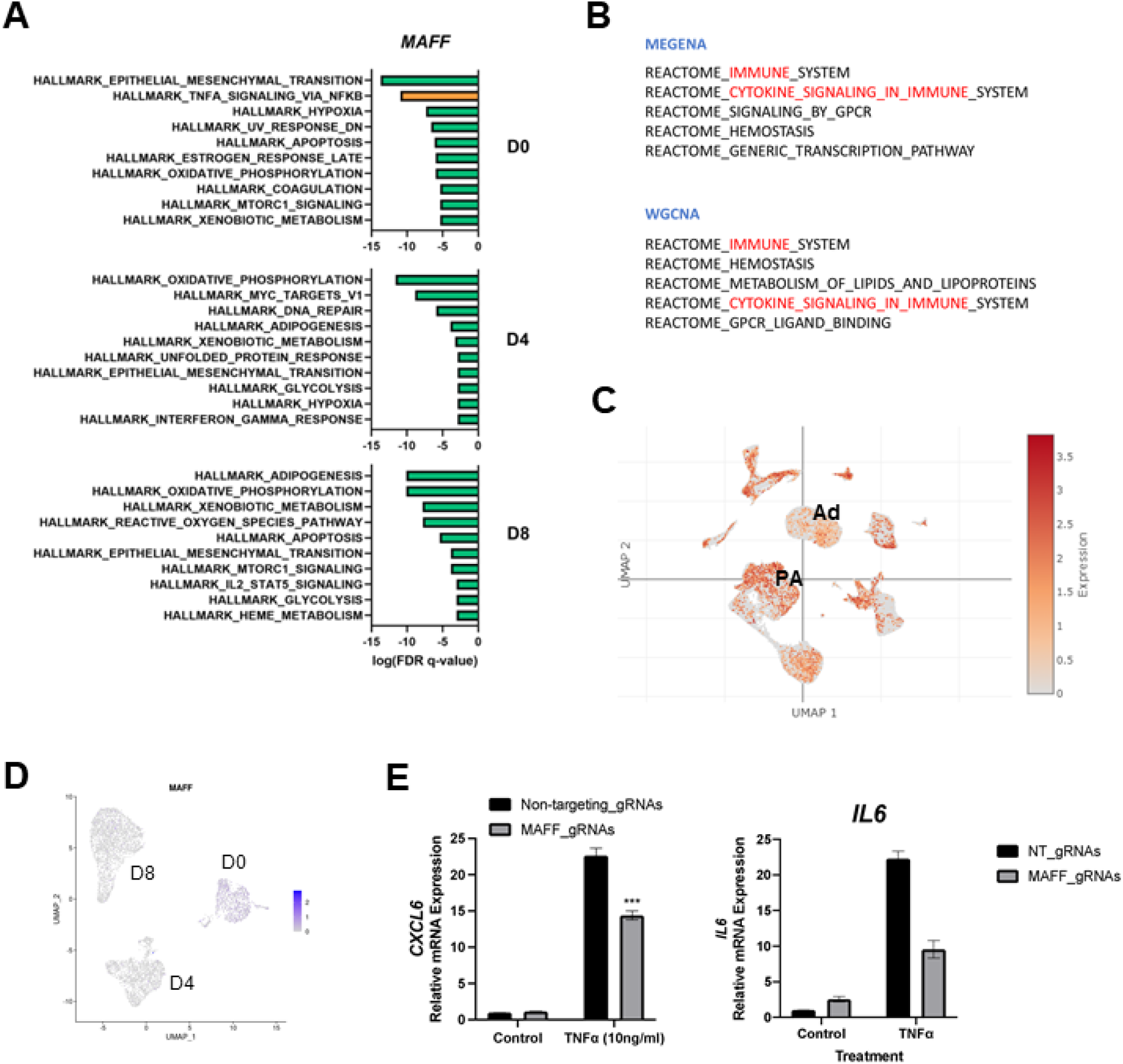
CROP-Seq identifies TNF-α gene expression response as a downstream target of MAFF. **A)** The results of Gene Set Enrichment Analysis for CROP-Seq results in *MAFF* knock-down cells at D0, D4 and D8 of differentiation. Gene sets relevant for TNF-α signaling is highlighted in orange. **B)** Co-expression modules identified for *MAFF* in human and mouse adipose tissue, constructed by WGCNA and MEGENA. The modules relevant for immune function are highlighted in red. **C)** UMAP plot showing single-cell RNA expression of *MAFF* in human adipose tissue^59^. Preadipocyte (PA) and adipocyte (Ad) clusters are marked. **D)** UMAP plot showing single-cell MAFF expression in SGBS cells in the CROP-Seq experiment. **E)** RT-qPCR quantification of *CXCL6* and *IL6* gene expression in *MAFF*-deficient preadipocytes than in non-targeting control preadipocytes following stimulation with TNFα. Two-tailed Student t Test. ***, p<0.001. N=3 biological replicates.

In addition to the impact of *MAFF* knock-down on disruption of gene expression networks at D0, *MAFF* has high expression in preadipocytes compared to adipocytes in human adipose tissue **(Figure 3C)** and is predominantly expressed in preadipocytes in the SGBS cell model **(Figure 3D)**. Therefore, for functional follow-up studies we focused on the hypothesis that *MAFF* plays a role in regulating immune function in preadipocytes. To test this hypothesis, we created individual SGBS preadipocyte lines with *MAFF* knock-down along with a control line with non-targeting sgRNAs, and subjected them to stimulation with TNF-α. Under *MAFF* knock-down, SGBS cells showed a blunted response to TNF-α treatment, quantified by the lower induction of the expression of genes encoding proinflammatory cytokines CXCL6 and IL-6 **(Figure 3E)**. We concluded that MAFF is a positive regulator of cytokine production in response to TNF-α in SGBS preadipocytes. In addition, TNF-α is a known negative regulator of adipogenesis^47^, and the changes in intracellular signaling in response to this cytokine under *MAFF* knock-down may directly impact adipogenesis, which was affected by *MAFF* knock-down at D4 and D8 **(Figure 3A)**.

### 3.5 *TIPARP* knock-down induces the expression of browning markers in differentiating SGBS cells

Next, we focused on another candidate gene, *TIPARP*, which was identified in a GWAS on WHRadjBMI^48^ and HDL and had an eQTL in subcutaneous adipose tissue. Gene Set Enrichment Analysis of *TIPARP* knock-down showed an unexpected enrichment of DE genes within pathways related to myogenesis at D4 and D8, and oxidative phosphorylation at D0 and D4 **(Figure 4A)**. Moreover, co-expression analysis in adipose tissue indicated potential TIPARP involvement in mitochondrial function **(Figure 4B)**. White adipocytes can undergo beiging^4^, a process in which their morphology and molecular profile becomes similar to brown adipocytes that develop from myogenic progenitors^49^. Brown and beige adipocytes contain significantly more mitochondria than white adipocytes, and they express uncoupling protein 1 (*UCP1*) which leads to the uncoupling of respiration from ATP production^4^. As the Hallmark database does contain few gene sets pertaining to mitochondria, we examined the enrichment of DE genes in *TIPARP* knock-down cells at D0 using Gene Ontology sets. Indeed, the most significant association within the Cellular Component gene sets was for mitochondria **(Supplementary Figure 4)**.

**Figure 4:**
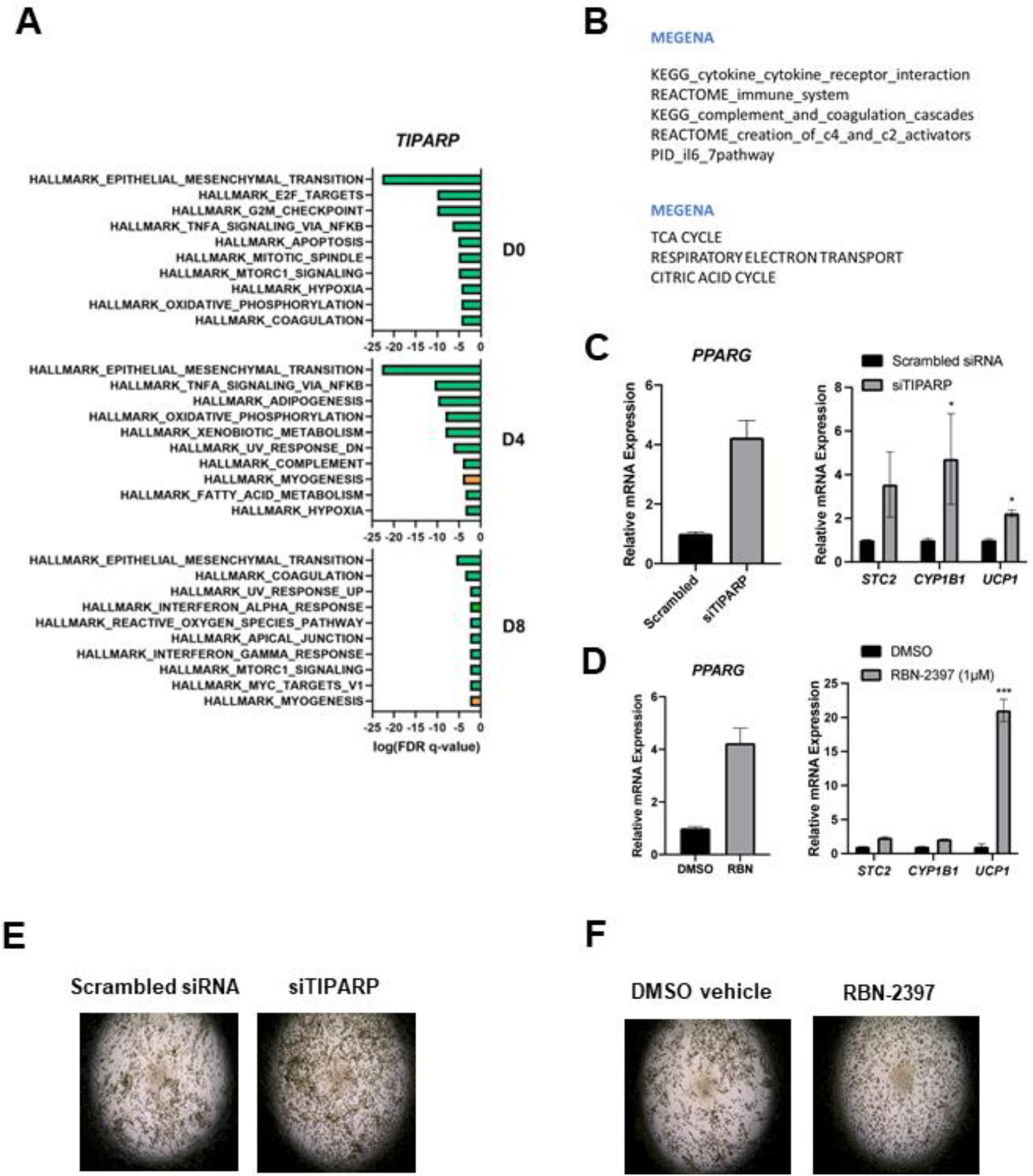
CROP-seq as an approach to connect changes in transcriptome to gene activity. **A)** The results of Gene Set Enrichment Analysis for CROP-Seq results in *TIPARP* knock-down cells at D0, D4 and D8 of differentiation. **B)** Co-expression network of *TIPARP* in adipose tissue. **C)** RT-qPCR quantification of differentiation marker PPARG and brown/beige adipocyte markers (STC2, CYP1B1, UCP1) gene expression in SGBS cells differentiated following gene knock-down using *TIPARP* siRNA (siTIPARP) or scrambled control siRNA. **D)** RT-qPCR quantification of PPARG, STC2, CYP1B1, and UCP1 in SGBS cells differentiated in the presence of TIPARP inhibitor RBN-2397 or DMSO control. **E)** Representative light microscopy images of SGBS cells differentiated following gene knock-down using *TIPARP* siRNA (siTIPARP) or scrambled control siRNA. **F)** Representative light microscopy images of SGBS cells differentiated in the presence of TIPARP inhibitor RBN-2397 or DMSO control. Two-tailed Student t test. ***, p<0.001; *, p<0.05. N=3 biological replicates.

To determine if there is an indication of beiging in *TIPARP* knock-down adipocytes, we compared the expression of brown/beige adipocyte marker *UCP1* and adipocyte marker *PPARG* following acute siRNA-mediated *TIPARP* knock-down in SGBS cells which then underwent differentiation. Both *UCP1* and *PPARG* had increased expression levels in *TIPARP* knock-down cells compared to control **(Figure 4C)**. Next, we validated these findings using specific TIPARP inhibitor RBN-2397^50^ **(Figure 4D)**. Both *TIPARP* knock-down and chemical inhibition led to increased and homogenous differentiation of small adipocytes **(Figure 4E,F)**. We conclude that TIPARP is a negative regulator of adipocyte beiging.

## 4. Discussion

To derive biological knowledge from associations between human genetic loci and trait/disease risk, functional testing is indispensable. Unbiased approaches to functional testing of candidate genes allow for the discovery of unexpected mechanisms. Here, we established the CROP-Seq platform for parallel gene expression manipulation using CRISPRi, with single cell transcriptome readout in human preadipocytes and adipocytes. Based on the results of the disruption of known regulators of adipogenesis, this approach allows for reliable identification of key pathways that are dysregulated following target gene downregulation. Moreover, the CROP-Seq approach can be used as an efficient tool for hypothesis generation regarding the biological function of disease-associated candidate genes in adipogenesis, preadipocyte and adipocyte function. This functional genomics approach significantly speeds up the translation of genetic associations to the knowledge of gene function. Therefore, CROP-Seq has the potential to bridge the current gap between hundreds of identified genomic associations and our limited understanding of molecular mechanisms of metabolic disease.

Beyond recapitulating the genes and pathways underlying the known regulators of adipocyte biology *CEBPB* and *PPARG*, we also uncovered biological pathways and phenotypes resulting from the knock-down of two candidate genes *MAFF* and *TIPARP*. Based on the analysis of cells with *MAFF* knock-down and functional validation tests, we identified *MAFF* as a novel regulator of transcriptional response to TNF-α in preadipocytes. *In vivo*, preadipocytes are present within adipose tissue alongside blood vessels^51^, and they are known to participate in inflammation^52^. In addition, the differentiation ability of human preadipocytes is impaired by inflammatory signals, including TNF-α, which are more prevalent in obesity^53^. Therefore, differences in *MAFF* function between individuals may contribute to varied levels of cytokine production by preadipocytes in response to pro-inflammatory environment in obese adipose tissue, leading to differences in adipose tissue maintenance or function. In fact, our CROP-Seq analysis indicated changes in the expression of adipogenesis-related genes at D4 and D8 in *MAFF* knock-down cells. In the future, *in vivo* models should be used to identify the effect of *MAFF* loss in preadipocytes on other cell types within adipose tissue, both under standard conditions and in obesity.

We also investigated the role of the TCDD Inducible Poly(ADP-Ribose) Polymerase (*TIPARP*). We found that *TIPARP* downregulation and inhibition increased the expression of adipocyte beiging marker *UCP1* in adipocytes. *TIPARP* is a known repressor of the aryl hydrocarbon receptor (AHR) and mediates the effects of dioxin on steatohepatitis^54^. However, recently an additional function of *TIPARP* was described as a repressor of several transcription factors, including HIF-1, c-Myc, and estrogen receptor^55^. Moreover, the specific TIPARP inhibitor RBN-2397 is undergoing a clinical trial as a drug capable of restoring type I interferon response in cancer cells^56^. The molecular pathway through which TIPARP affects adipogenesis or adipocyte function remains to be elucidated.

CROP-Seq has several drawbacks due to the inclusion of scRNA-Seq as the readout. scRNA-Seq is not well suited for detecting expression changes for genes with low baseline expression level. To allow for the verification of target gene knock-down, only genes with a medium to high expression should be targeted. To avoid the expression limitation, other methods such as Targeted Perturb-Seq (TAP-Seq)^57^ have been developed. In addition, pooled gene knock-down inevitably leads to the presence of cells with more than one sgRNA present. scMAGeCK^17^ and criprQTL^58^ pipelines were developed to include cells with multiple sgRNAs, which need more validation and improvement.

The pooled approach used in the CROP-Seq leads to additional limitations with regards to studying biological effects of genes encoding secreted factors (e.g. *ADIPOQ* and *PDGFC*) because cells with target gene knock-down might still be exposed to the factors if they are secreted by other cells in their environment. In our CROP-Seq experiment we observed robust changes in gene expression in response to knock-down of genes encoding transcription factors (*CEBPB, PPARG, MAFF*) and an enzyme (*TIPARP*). Therefore, we envision CROP-Seq as applicable to study genes encoding a variety of protein types, although it is likely that the interpretation of the results is more straightforward if the candidate proteins directly affect gene expression that is quantified by CROP-Seq.

Adipocytes, both *in vivo* and *in vitro*, are technically challenging to work with, because of their large size, high lipid content, and fragility. To ensure that adipocytes are well-represented within the sequenced cells from the differentiated population, we propose that cell sorting based on fluorescent staining of neutral lipids or cell size and granularity can be added to the CROP-Seq pipeline.

In summary, our method is the first to combine pooled single-cell CRISPRi knock-down with scRNA-Seq readout in a human adipogenesis model. The CROP-Seq is a powerful tool to investigate molecular function of novel genes with putative role in adipocyte biology.

## Abbreviations

ATP: adenosine triphosphate
CRISPRi: Clustered Regularly Interspaced Short Palindromic Repeats interference
KRAB-dCas9: Krüppel-associated box repressor – inactive CRISPR associated protein 9
scRNA-Seq: single-cell RNA sequencing
SGBS: Simpson-Golabi-Behmel syndrome

## Author contributions

The experiments were designed by J.L., C.Y.P., T.Q. and J.W.K. and executed by J.L. Computational analyses were conducted by D.S., M.B., M.J.G., and X.Y. Additional technical expertise or material support was provided by P.S.G., I.C.-O., C.Y.P., M.W., and K.J.S. The manuscript was written by E.B.-M. with input from all the authors.

## Acknowledgements

We would like to thank Erik Ingelsson and the members of Knowles and Quertermous labs for their feedback on the project. We acknowledge the technical assistance of the Stanford Genomics Service Center and the Stanford Shared FACS Facility.

