## Supplementary data for "CROP-Seq: a single-cell CRISPRi platform for characterizing candidate genes relevant to metabolic disorders in human adipocytes"

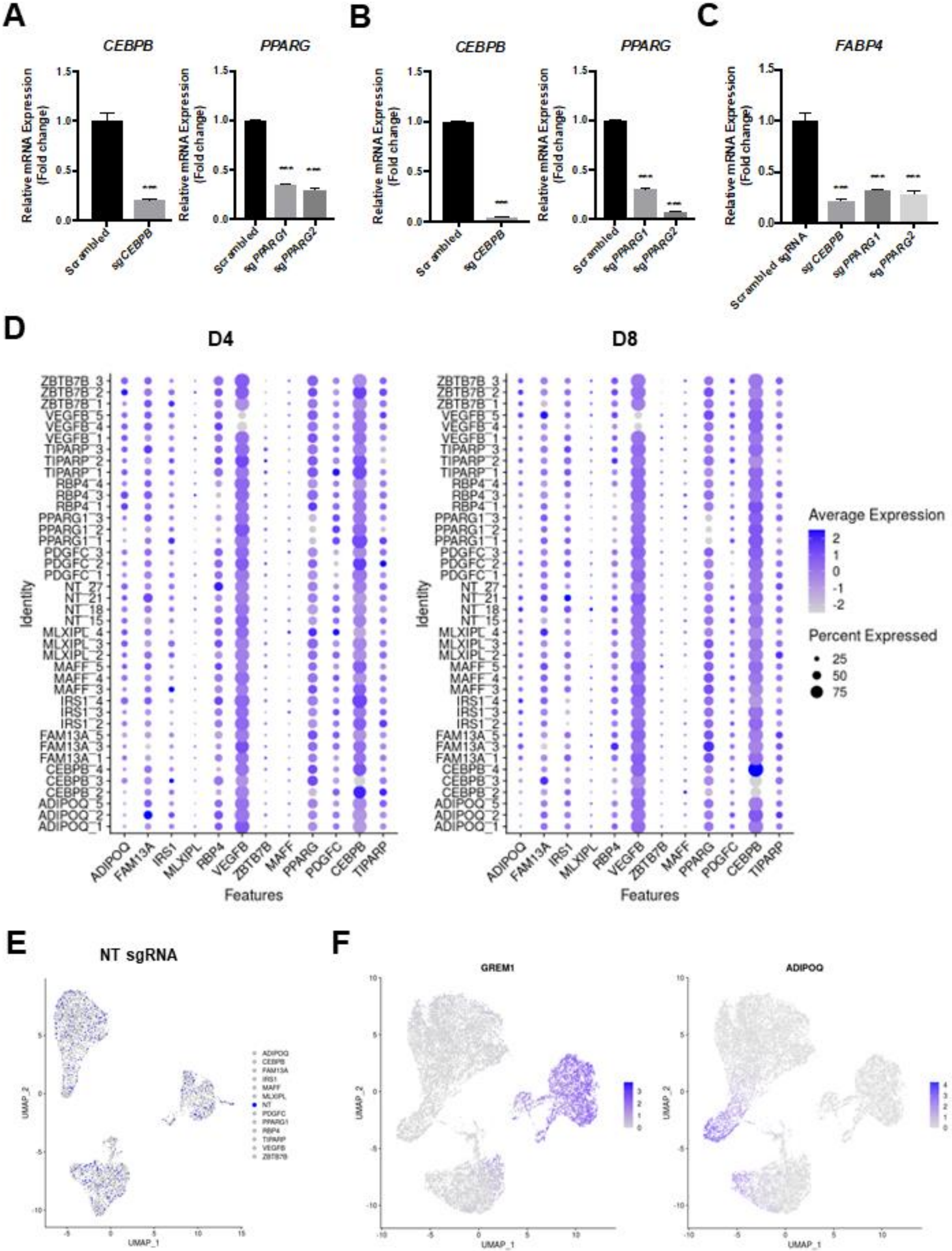

A
CEBPB
PPARG
Relative mRNA Expression (Fold change)
Scrambled
sgCEBPB
sgPPARG1
sgPPARG2
***
B
CEBPB
PPARG
C
FABP4
Scrambled sgRNA
D
D4
D8
Average Expression
Percent Expressed
25
50
75
Identity
Features
ADIPOQ
FAM13A
IRS1
MLXIPL
RBP4
VEGFB
ZBTB7B
MAFF
PPARG
PDGFC
CEBPB
TIPARP
ZBTB7B_3
ZBTB7B_2
ZBTB7B_1
VEGFB_5
VEGFB_4
VEGFB_1
TIPARP_3
TIPARP_2
TIPARP_1
RBP4_4
RBP4_3
RBP4_1
PPARG1_3
PPARG1_2
PPARG1_1
PDGFC_3
PDGFC_2
PDGFC_1
NT_27
NT_21
NT_18
NT_15
MLXIPL_4
MLXIPL_3
MLXIPL_2
MAFF_5
MAFF_4
MAFF_3
IRS1_4
IRS1_3
IRS1_2
FAM13A_5
FAM13A_3
FAM13A_1
CEBPB_4
CEBPB_3
CEBPB_2
ADIPOQ_5
ADIPOQ_2
ADIPOQ_1
E
NT sgRNA
UMAP_1
UMAP_2
NT
PPARG1
F
GREM1
ADIPOQ

**Supplementary Figure 1**: Supplementary data for Figure 1.

**A)** RT-qPCR analysis of *CEBPB* and *PPARG* expression in SGBS at D0 of differentiation; dCas9-BFP-KRAB cells transduced with either a non-targeting sgRNA (scrambled) or 3 pooled sgRNAs against *CEBPB*, *PPARG1* and *PPARG2*, respectively.

**B)** RT-qPCR analysis of *CEBPB* and *PPARG* expression in SGBS at D8 of differentiation; dCas9-BFP-KRAB cells transduced with either a non-targeting sgRNA (scrambled) or 3 pooled sgRNAs against *CEBPB*, *PPARG1* and *PPARG2*, respectively.

**C)** RT-qPCR analysis of the differentiation marker, *FABP4*, in SGBS at D8 of differentiation; dCas9-BFP-KRAB adipocytes transduced with either a non-targeting sgRNA or 3 pooled sgRNAs against *CEBPB*, *PPARG1* and *PPARG2*, respectively.

**D)** Quantification of target gene expression at D4 and D8 by average expression and percent of cells with detectable expression depending on the sgRNA.

**E)** UMAP plot of all cells showing clustering of cells expressing non-targeting gRNAs (NT sgRNA).

**F)** UMAP plot of all cells showing single-cell expression of marker genes for pre-adipocytes (*GREM1*) and adipocytes (*ADIPOQ*).

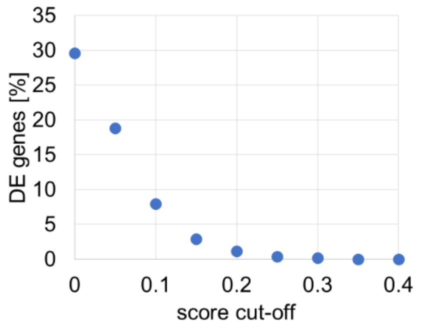


**Supplementary Figure 2**: Percentage of genes identified as DE in *PPARG* knock-down at D8, depending on the score cut-off used on the list of genes identified using scMAGeCK pipeline. An additional cutoff of p-value < 0.05 was kept constant throughout the analysis.

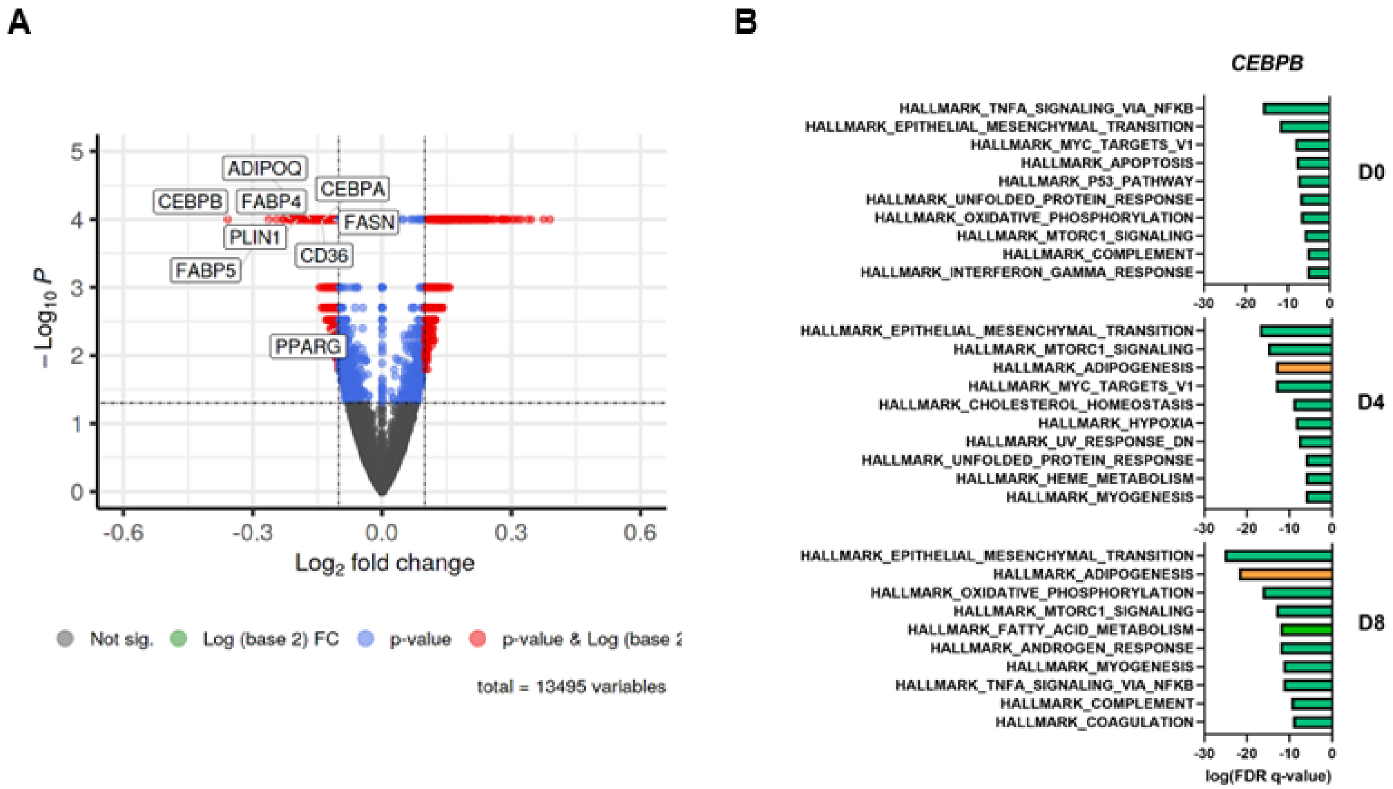


**Supplementary Figure 3:** Supplementary data for Figure 2.

- **A)** Volcano plot of DE genes for *CEBPB* knock-down at D0.
- **B)** GSEA analysis of *CEBPB* knock-down DE genes at D0, D4, D8. Adipogenesis highlighted in orange.

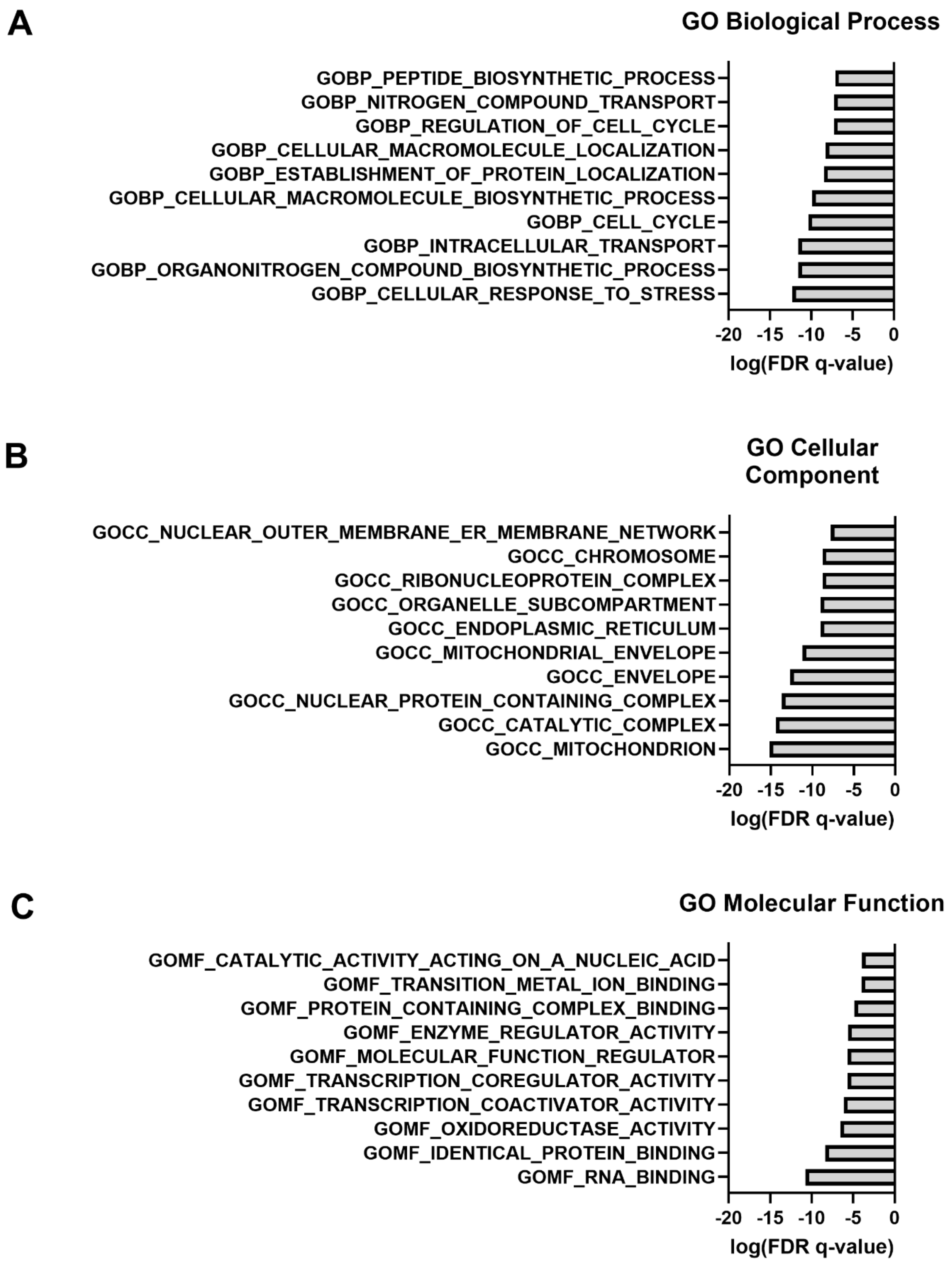


**Supplementary Figure 4:** Supplementary data for Figure 4. GSEA for *TIPARP* knock-down at D0 using Gene Ontology datasets. Top 10 datasets for each subset are shown.

A) GSEA analysis for Gene Ontology: Biological Process.
B) GSEA analysis for Gene Ontology: Cellular Component.
C) GSEA analysis for Gene Ontology: Molecular Function.

| sgRNA # | sgRNA name | Target gene | sgRNA sequence |
|---|---|---|---|
| 1 | ADIPOQ_1 | *ADIPOQ* | GTTGCTCTTACCCTCTGGTA |
| 2 | ADIPOQ_2 | | GGCTTGACCAGGTAGGGCAA |
| 3 | ADIPOQ_5 | | GCCACAGACTGCAGTCAGAA |
| 4 | FAM13A_1 | *FAM13A* | GACGCTTTCTGAGAGAATGG |
| 5 | FAM13A_3 | | GTCCGCTGAACCCACATGGC |
| 6 | FAM13A_5 | | GTAGCCATCTGTGTAAGTAT |
| 7 | IRS1_2 | *IRS1* | GTCTCCTCCGAGTGAGACGG |
| 8 | IRS1_3 | | GACCGCGCCGCCGTCTCACT |
| 9 | IRS1_4 | | GGAGCCCCAACCAAAACAAG |
| 10 | MLXIPL_2 | *MLXIPL* | GAGGGACCAGGCGGTTGCGG |
| 11 | MLXIPL_3 | | GCTCCGGGAGCCGCAGGACC |
| 12 | MLXIPL_4 | | GGCAGGACCGGGCTCGGGTG |
| 13 | RBP4_1 | *RBP4* | GCCCGGCCGCGACTCACCAC |
| 14 | RBP4_3 | | GCGCGCGGTTCCCCTCCCGG |
| 15 | RBP4_4 | | GAAGCCTGGCCGCCGAGTCC |
| 16 | VEGFB_1 | *VEGFB* | GGGCGCGTGCATCCCCTGGG |
| 17 | VEGFB_4 | | GCCCGGCGCGTGCATCCCCT |
| 18 | VEGFB_5 | | GCCTCGGCCGAGGGGATCTG |
| 19 | ZBTB7B_1 | *ZBTB7B* | GGGAGCCCCCCAGCAGCGAG |
| 20 | ZBTB7B_2 | | GCTGCTGAGGCTCACAGTGT |
| 21 | ZBTB7B_3 | | GGCGGCCTTCTGACCAGGAC |
| 22 | MAFF_3 | *MAFF* | GTGCTCCTACGTGACTCGGT |
| 23 | MAFF_4 | | GGCCCGGGCCCGATCCGTCG |
| 24 | MAFF_5 | | GGCGGCGGCGCCGGGAGGAA |
| 25 | PPARG1_1 | *PPARG* | GGAGTGAGCCAACTCACCTA |
| 26 | PPARG1_2 | | GGCCAACTCACCTAAGGAAA |
| 27 | PPARG1_3 | | GTCAGTGGTAGGTAAGGAAG |
| 28 | PDGFC_1 | *PDGFC* | GAGACTGGAAGCCAAGTCGG |
| 29 | PDGFC_2 | | GGACAGAGGCGAAAACTCAA |
| 30 | PDGFC_3 | | GCCCGGAGAGCCGCATCTAT |
| 31 | NT_15 | Non-targeting | GGCACAGCATCAGTACCCGA |
| 32 | NT_18 | | GGGTAGATAGCACCGCTTAC |
| 33 | NT_27 | | GAGGCGGCGCTAACCTCGGG |
| 34 | NT_21 | | GGAGTGCTGAACCACTGCGT |
| 35 | CEBPB_2 | *CEBPB* | GGCCCAGCGGCTCCAGGTAC |
| 36 | CEBPB_3 | | GTAGTCGTCGGAGAAGAGGT |
| 37 | CEBPB_4 | | GTGGTGGCTGGACCCTCGGG |
| 38 | TIPARP_1 | *TIPARP* | GGCTCCGGGAGGGCACTGAG |
| 39 | TIPARP_2 | | GGGACCACAGAGCCGCGACG |
| 40 | TIPARP_3 | | GCTTAGGCACTTAGAGCCTG |

**Supplementary Table 1:** Sequences of sgRNAs used in the CROP-Seq screen.
